# Efficient repositioning of approved drugs as anti-HIV agents using Anti-HIV-Predictor

**DOI:** 10.1101/087445

**Authors:** Shao-Xing Dai, Huan Chen, Wen-Xing Li, Yi-Cheng Guo, Jia-Qian Liu, Jun-Juan Zheng, Qian Wang, Hui-Juan Li, Bi-Wen Chen, Yue-Dong Gao, Gong-Hua Li, Yong-Tang Zheng, Jing-Fei Huang

**Author notes:** These authors contributed equally to this work (Shao-Xing Dai, Huan Chen, Wen-Xing Li). Corresponding author: Shao-Xing Dai Gong-Hua Li Yong-Tang Zheng Jing-Fei Huang.

## Abstract

Development of new, effective and affordable drugs against HIV is urgently needed. In this study, we developed a world’s first web server called Anti-HIV-Predictor (http://bsb.kiz.ac.cn:70/hivpre) for predicting anti-HIV activity of given compounds. This server is rapid and accurate (accuracy >93% and AUC > 0.958). We applied the server to screen 1835 approved drugs for anti-HIV therapy. Totally 67 drugs were predicted to have anti-HIV activity, 25 of which are anti-HIV drugs. Then we experimentally evaluated 35 predicted new anti-HIV compounds by assays of syncytia formation, p24 quantification, cytotoxicity. Finally, we repurposed 7 approved drugs (cetrorelix, dalbavancin, daunorubicin, doxorubicin, epirubicin, idarubicin and valrubicin) as new anti-HIV agents. The original indication of these drugs is involved in a variety of diseases such as female infertility and cancer. Anti-HIV-Predictor and the 7 repurposed anti-HIV agents provided here demonstrate the efficacy of this strategy for discovery of new anti-HIV agents.

## Introduction

Even 30 years after its discovery, human immunodeficiency virus (HIV) remains a great threat to humans^1,2^. Acquired immune deficiency syndrome(AIDS), the disease elicited by HIV infection, is considered to be pandemic and represents the greatest global public health crisis^3^. There are an estimated 39 million deaths caused by AIDS since its first recognition^4^. According to the report of the World Health Organization(WHO), there were approximately 37 million people living with HIV at the end of 2014 with 2 million people becoming newly infected with HIV in 2014 globally. And 1.2 million people died from HIV-related causes globally in 2014 (http://www.who.int/mediacentre/factsheets/fs360/en/).

Many scientists around the world are committed to finding scientifically proven strategies for HIV prevention and treatment. Recent decades, significant progress has been achieved in the development of vaccines and drugs against HIV infection. Several clinical trials of anti-HIV vaccines, including RV144, are ongoing^5,6^. The RV144 trial demonstrated 31% vaccine efficacy at preventing human immunodeficiency virus (HIV)-1 (referred to as HIV for the rest of this study) infection^7^. The most notable achievement is the transformation of HIV/AIDS from an inevitable death sentence to a chronic illness by the introduction of combination antiretroviral therapy^8,9^. More than thirty anti-HIV drugs have been approved by the US Food and Drug Administration (FDA)^10^. These drugs act mainly on reverse transcriptase, protease, integrase, CCR5 and so on^11^. Behind the progress, many studies were carried out to discover anti-HIV drug candidates by screening a large number of natural or synthetic compounds. A representative study was the AIDS anti-viral screen program of the National Cancer Institute (NCI), which screened more than 30,000 compounds (https://dtp.cancer.gov/)^12,13^. After this program, many anti-HIV compounds were reported and deposited in ChEMBL database (ChEMBLdb)^14^. These data are helpful for data mining and developing new tool toward HIV treatment. Despite considerable progress, treatment of AIDS still faces multiple challenges^15,16^. To date, no truly effective drug able to eliminate HIV has been developed^17^. Furthermore, HIV is highly variable and can quickly acquire resistance against any drug with which it is confronted^11,18^. Therefore, there is a constant demand to develop new, effective and affordable anti-HIV drugs. In the past decades, tens of millions of chemical compounds have been deposited in public databases^19^. Screening these huge databases for new anti-HIV drugs through experimental methods is a tedious, expensive and time-consuming process. The time and money-saving way is that all compounds in the database are firstly filtered by the computational analysis of the anti-HIV potential, then evaluated by experiment. Therefore, a rapid and accurate computational method is urgently required for predicting anti-HIV activity of chemical compounds.

In this study, we aim to establish a web server to predict anti-HIV activity of given compound and apply the web server to discover new anti-HIV agents through drug repositioning of FDA approved drugs (Figure 1). Drug repositioning is the process of finding new uses outside the scope of the original medical indication for existing drugs^20,21^. An advantage of drug repositioning lies in the fact that the safety, dosage, and toxicity of existing drugs have already been vetted^22^. Therefore, repurposed candidate drugs can often enter clinical trials much more rapidly than newly developed drugs^23^. Recently, many anti-infectious agents have been discovered to combat pathogens using drug repositioning^24–28^.

**Figure 1.**
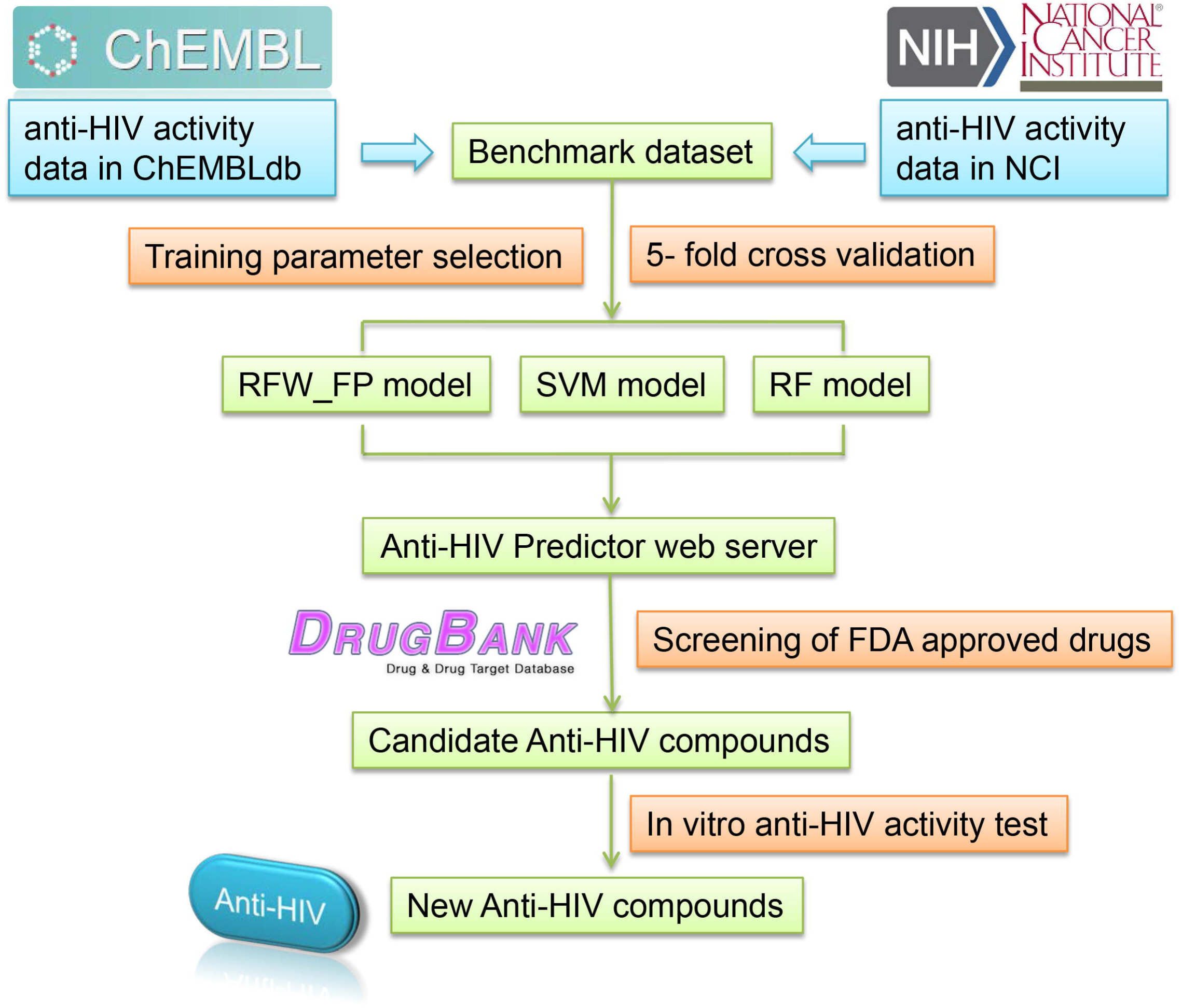
The flowchart of Anti-HIV-Predictor and drug repositioning. After construction of benchmark dataset, three models (RFW_FP model, SVM model and RF model) were generated to predict anti-HIV activity of chemical compounds by training, parameter selection and 5-fold cross validation. The web server Anti-HIV-Predictor was established by incorporating the three prediction models. The web server was used to screen all FDA approved drugs. Finally, the predicted new anti-HIV compounds were evaluated for anti-HIV activity *in vitro*.

Therefore, in this study, we firstly developed three rapid and accurate computational methods to predict anti-HIV activity of a given compound. Then a web server called Anti-HIV-Predictor (http://bsb.kiz.ac.cn:70/hivpre) is established by integrating the three methods. This web server is free and open to all users. All FDA approved drugs were screened using the web server. Finally, the predicted new anti-HIV compounds were selected for *in vitro* testing of anti-HIV activity. Using this strategy, we identified cetrorelix, dalbavancin and five anthracycline drugs as new potent anti-HIV agents.

## Results

### Development of Anti-HIV-Predictor

Workflow for establishing Anti-HIV-Predictor is outlined in Figure 1. Anti-HIV-Predictor firstly integrated all the data of anti-HIV activity from ChEMBL and NCI database to construct benchmark dataset. Then, using the benchmark dataset, three prediction models were generated by training, parameter selection and validation. The first model is relative frequency-weighted fingerprint (RFW_FP) based model. RFW_FP is a novel molecular description method which considers the frequency of bit in active and inactive datasets and integrates it to each compound fingerprint. RFW_FP was first used in our previous study and powerful to distinguish the active and inactive compounds for anti-cancer^29,30^. The other two models are Support Vector Machine (SVM) and Random Forest (RF) models. Last, three models (RFW_FP model, SVM model and RF model) were incorporated to predict anti-HIV activity of chemical compounds. The details for development of Anti-HIV-Predictor are given in the **Materials and methods** section.

### Performance of Anti-HIV-Predictor

The overall performance of the RFW_FP, SVM and RF models was quantified by receiver operating characteristic curve (ROC). For each model, the ROC was plotted and the area under the curve (AUC) was calculated (Figure 2a). The ROC curve shows the relation between true positive rate and false positive rate for each threshold of the real-value outputs. The AUC value of the RFW_FP, SVM and RF models are 0.958, 0.974 and 0.977, respectively. All three models achieve AUC value greater than 0.958, which reveals the excellent effectiveness of the models. From the three curves, we can also observe that the three models can effectively identify active anti-HIV compounds with high true-positive rates against low false positive rates.

**Figure 2.**
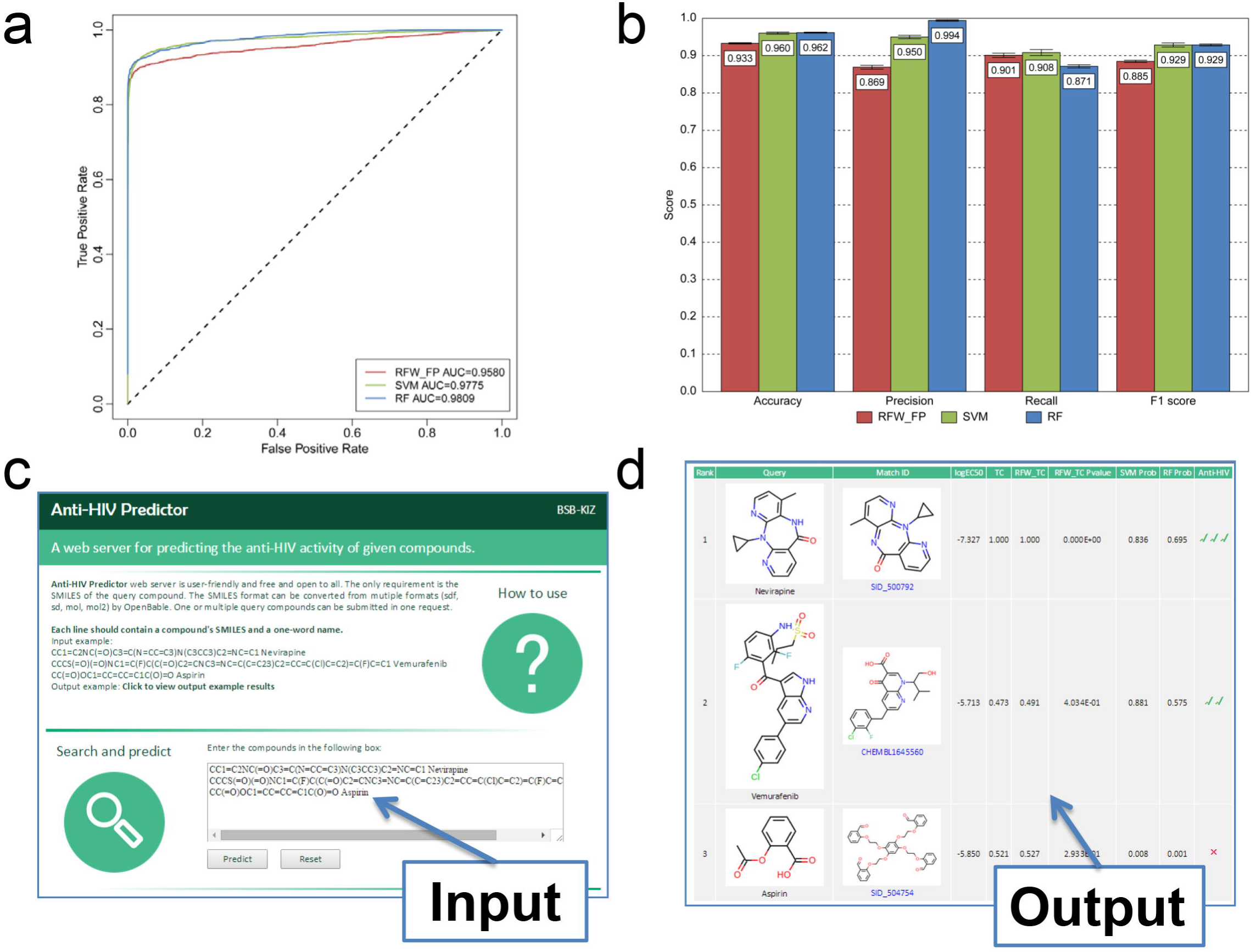
The performance, input and output of Anti-HIV-Predictor. **(a)** The ROC and AUC for the RFW_FP model (red), SVM model (green) and RF model (blue), respectively. **(b)** The statistical average results for 10 runs of 5-fold cross validation. The panel indicate the mean and standard deviation values of accuracy, precision, recall and F1 score derived from the RFW_FP model (red), SVM model (green) and RF model (blue), respectively. Vertical lines indicate the standard deviations (SDs). **(c)** Input interface of Anti-HIV-Predictor. The web server only needs the SMILES of the query compound as input. **(d)** The output of Anti-HIV-Predictor. The output contains the matched similar compound, the predicting information and the predicting conclusion whether the query compound has anti-HIV activity (see text for details). For example, Anti-HIV-Predictor assigns three ticks for the drug nevirapine and a cross for aspirin.

The classification performance of the models was also assessed in terms of accuracy, precision, recall and F1 score (Figure 2b). As 10 runs of 5-fold cross-validation (CV) method were used, these scores were averaged. Over the ten runs, their standard deviations were also reported. As shown in Figure 2b, the RFW_FP model obtains the statistical average of 93.3%, 86.9%, 90.1%, and 88.5% for accuracy, precision, recall, and F1 score, respectively. The accuracy, precision, recall, and F1 score of SVM model are 96%, 95%, 90.8%, and 92.9%, respectively. RF model performs best with the accuracy of 96% and precision of 99.4%.

### Input and output of Anti-HIV-Predictor

Anti-HIV-Predictor is user-friendly and free and open to all users. The only requirement of Anti-HIV-Predictor is the SMILES of the query compound. One or multiple query compounds can be submitted in one request (Figure 2c). The total number of input compounds is limited to 100 for each submission. Anti-HIV-Predictor needs about 60 seconds to load the background data and trained models required for prediction. Therefore, 1-10 compounds requires about 90 seconds, but 100 compounds only requires about 150 seconds. A query with 1–10 compounds requires about 90 seconds, whereas a query with 100 compounds only requires about 150 seconds. After calculated, the output of Anti-HIV-Predictor was shown in Figure 2d. Firstly, the output gives the most similar compound of the query compound. The structures, database links and anti-HIV activities (logEC_50_) of the matched similar compound were also displayed. Secondly, the output contains some important predicting information, for example, Tanimoto Coefficient score (TC), the Relative Frequency-Weighted Tanimoto Coefficient (RFW_TC), P-value of RFW_TC model, probability estimation by SVM model and RF model. Finally, the output shows the predicting conclusion whether the query compound has anti-HIV activity. One tick represents the query compound is predicted as anti-HIV compound by one of the three models. One cross means that all the three models show no anti-HIV activity for the query compound.

### Rapid and accurate computational screen of FDA approved drugs using Anti-HIV-Predictor

To discover new anti-HIV agents through drug repositioning of FDA approved drugs, 1835 approved drugs with SMILES string were downloaded from DrugBank (http://www.drugbank.ca). Using Anti-HIV-Predictor, all the drugs were screened rapidly by the three models. The results of computational screen are shown in Figure 3. Most drugs have no anti-HIV activity based on the prediction. These drugs were shown as blue dots with RFW_TC P-value ≥0.05, SVM probability and RF probability ≤0.5 (Figure 3a). The green dots represent the drugs with anti-HIV activity supported by one or two models (RFW_TC P-value <0.05 or SVM probability >0.5 or RF probability >0.5). The red dots represent the drugs with anti-HIV activity supported by all three models (RFW_TC P-value <0.05 and SVM probability >0.5 and RF probability >0.5). As shown in Figure 3b, totally 67 drugs were predicted to have anti-HIV activity by all three models. The RFW_FP, SVM and RF models predicted 240, 178, 110 drugs with anti-HIV activity, respectively. Therefore, the 67 drugs represent the intersection of the results of the three different models (Figure 3b). Among the 67 drugs, there are 25 approved anti-HIV drugs and 7 drugs with anti-HIV activity. For other 35 drugs, there is no experimental test for anti-HIV activity (**Table S1**).

**Figure 3.**
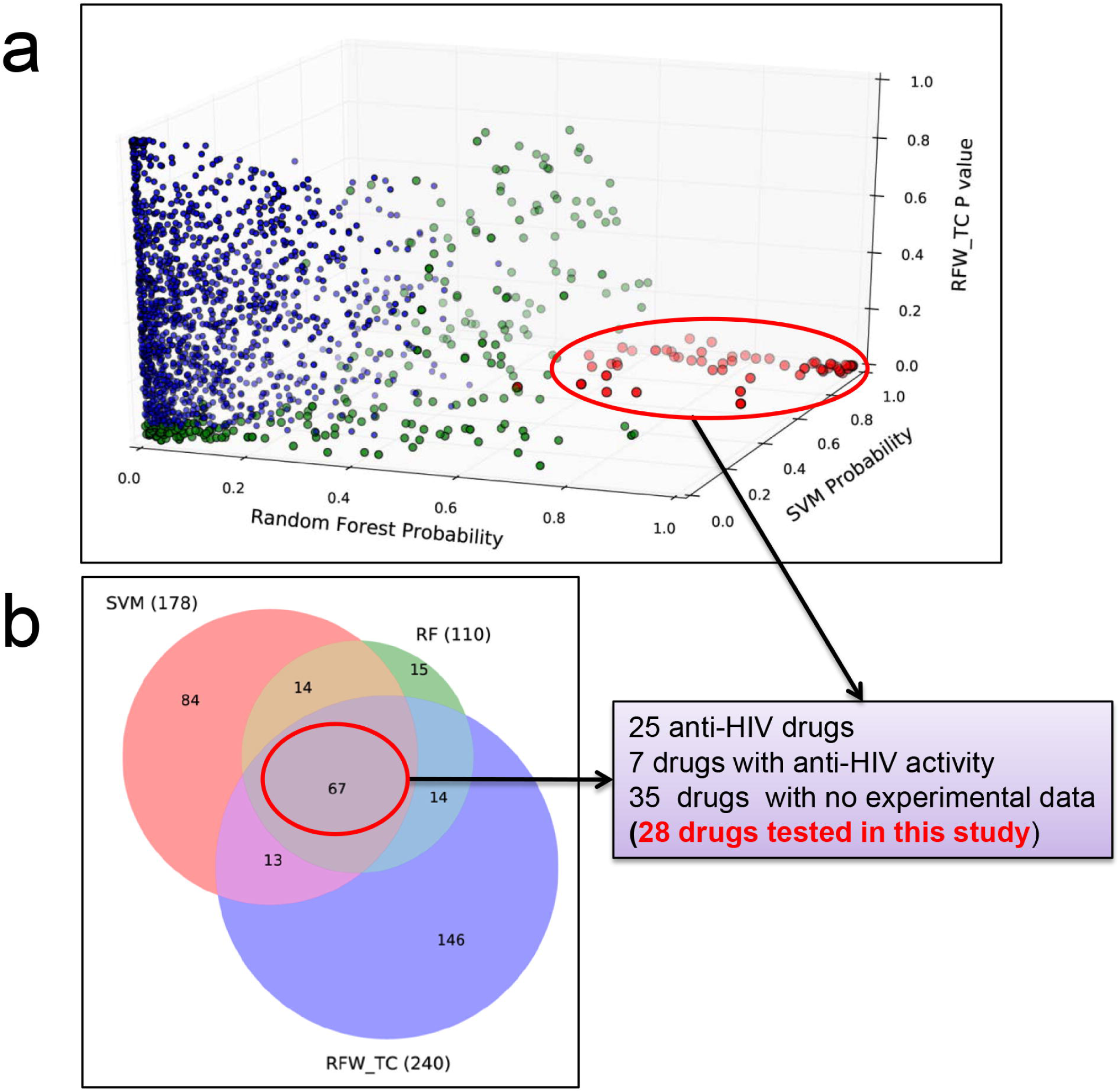
The results of computational screen of FDA approved drugs using Anti-HIV-Predictor. **(a)** Three-axis plot of all approved drugs based on the predict scores of the three models (RFW_FP model, SVM model and RF model). Each dot represents a drug. The blue dot means the drug with no anti-HIV activity. The green dot means the drug with anti-HIV activity supported by one or two models. The red dot indicates the drug with anti-HIV activity predicted by all three models. **(b)** Venn diagram of the screening results. The RFW_FP model, SVM model and RF model predicted 240, 178 and 110 anti-HIV drugs, respectively. The overlap is 67 drugs which are categorized into three groups: approved anti-HIV drugs (25), drugs with anti-HIV activity(7) and drugs with no experimental data (35).

### Experimental confirmation of 15 approved drugs with anti-HIV activity

As the 35 drugs have not been experimental test for anti-HIV activity, it is interesting and worth evaluating their anti-HIV activity by experiment. 28 of these drugs were purchased from CASMART (http://www.casmart.com.cn). Other 7 drugs are not purchased and tested because they are not available or very expensive. Therefore, a total of 28 drugs were evaluated for their anti-HIV activity *in vitro* with azidothymidine (AZT) as a positive control. The cytotoxicity of these compounds on T cell line C8166 was assessed by MTT colorimetric assay, and 50% cytotoxicity concentration (CC_50_) was calculated. The inhibitory effect of compounds on HIV replication was measured by the syncytia formation assay and 50% effective concentration (EC_50_) was calculated as described previously. The assay results of the 28 compounds are presented in Table 1. For comparison, AZT, the first anti-HIV drug approved by FDA, was utilized as the reference compound. As shown in Table 1, 15 compounds show anti-HIV activity with the EC_50_ values ranging from 0.004 to 93.794 μM. More than half of the tested compounds (15/28) exhibit activity against HIV. It indicates that Anti-HIV-Predictor is a powerful tool for discovering anti-HIV compounds.

**Table 1.** The cytotoxicity (CC_50_), anti-HIV-1 activity on HIV-1_NL4-3_ strain (CPE EC_50_), and therapeutic index (TI) of the tested 28 compounds

| Drug name | Drug ID | CC<sub>50</sub> (μM)<br>Mean±SD | EC<sub>50</sub> (μM)<br>Mean±SD | TI |
| --- | --- | --- | --- | --- |
| Bivalirudin | DB00006 | >200 | >200 | Inactivity |
| Carbetocin | DB01282 | >200 | >200 | Inactivity |
| **Cetrorelix** | **DB00050** | **>200** | **15.285±1.024** | **>12.3<sup>a</sup>** |
| Cytarabine | DB00987 | 101.819±14.844 | 54.603±14.602 | 2.9-1.3 |
| **Dalbavancin** | **DB06219** | **>200** | **8.694±0.000** | **>23.0** |
| **Daunorubicin** | **DB00694** | **0.279±0.085** | **0.029±0.010** | **5.0-19.2** |
| Desmopressin | DB00035 | >200 | >200 | Inactivity |
| **Doxorubicin** | **DB00997** | **0.191±0.017** | **0.013±0.001** | **13.4-17.3** |
| **Epirubicin** | **DB00445** | **0.125±0.042** | **0.007±0.003** | **9.2-41.8** |
| Gatifloxacin | DB01044 | 116.743±24.863 | 126.263±16.005 | 0.6-1.3 |
| Gonadorelin | DB00644 | >200 | 72.417±3.403 | >2.6 |
| **Idarubicin** | **DB01177** | **0.077±0.004** | **0.004±0.001** | **18.3-26.7** |
| **Levofloxacin** | **DB01137** | **>200** | **17.533±1.456** | **>10.5** |
| Linaclotide | DB08890 | >200 | >200 | Inactivity |
| Moxifloxacin | DB00218 | 102.902±13.248 | 104.286±13.324 | 0.8-1.3 |
| Nafarelin | DB00666 | >200 | >200 | Inactivity |
| Ofloxacin | DB01165 | >200 | 114.097±12.820 | >1.6 |
| Pentagastrin | DB00183 | >200 | 93.794±42.202 | >1.5 |
| **Polymyxin B Sulfate** | **DB00781** | **109.157±0.879** | **15.007±4.186** | **5.6-10.2** |
| Sofosbuvir | DB08934 | >200 | >200 | Inactivity |
| Somatostatin | DB09099 | >200 | >200 | Inactivity |
| sparfloxacin | DB01208 | 85.112±11.066 | 31.594±2.029 | 2.2-3.3 |
| Terlipressin | DB02638 | >200 | >200 | Inactivity |
| Tolvaptan | DB06212 | 97.532±16.441 | 15.816±1.101 | 4.8-7.7 |
| Triptorelin | DB06825 | >200 | >200 | Inactivity |
| **Valrubicin** | **DB00385** | **2.405±0.446** | **0.082±0.002** | **23.3-35.6** |
| Vancomycin | DB00512 | >200 | >200 | Inactivity |
| Verteporfin | DB00460 | 7.502±3.177 | 10.235±4.213 | 0.3-1.8 |
| AZT | DB00006 | 1031.353±286.058 | 0.004±0.000 | >181324 |

a. The drugs with TI value more than 10 were highlighted with bold fonts.

### Identification of 7 approved drugs as new anti-HIV agents

Among the 15 compounds above, some compounds show serious cytotoxicity and then result in a very low therapeutic index (TI). The drugs with TI value more than 10 were further evaluated their anti-HIV activity by quantification of HIV p24 expression using ELISA method^31^. The best 7 drugs based on the results of cytotoxicity, syncytia formation and p24 quantification assays were displayed **in Table 2 and Figure 4.** Among the best 7 drugs, cetrorelix and dalbavancin are polypeptides, while other five drugs daunorubicin, doxorubicin, epirubicin, idarubicin and valrubicin belong to the class of anthracyclines. Cetrorelix, a synthetic decapeptide, is used for the inhibition of premature luteinizing hormone (LH) surges in women undergoing controlled ovarian stimulation^32^. Dalbavancin, a second-generation lipoglycopeptide antibiotic, is approved for the treatment of acute bacterial skin and skin structure infections caused by the gram-positive pathogens^33^. Cetrorelix and dalbavancin exhibit anti-HIV activity with EC_50_ of 1.788±0.115 and 1.296±0.186 μM, respectively. No cytotoxicity was detected for cetrorelix and dalbavancin. The cytotoxicity CC_50_ of cetrorelix and dalbavancin are both more than 200 μM. The percent viability at the concentration EC_50_ is almost 100% for cetrorelix and dalbavancin (Figure 4). Therefore, cetrorelix and dalbavancin show a very high therapeutic index (TI>105 and TI >135, respectively). The five anthracycline drugs are approved for the treatment of acute myeloid leukemia, bladder and breast cancer and so on ^34^. These anthracycline drugs show strong anti-HIV activity with EC_50_ varying from 0.003∼0.076 μM. The anti-HIV activity of Idarubicin is close to or better than that of AZT (0.003 μM for Idarubicin vs 0.005 μM for AZT). However, these anthracycline drugs exhibit a certain degree of cytotoxicity (Figure 4). The percent viability at the concentration EC_50_ is ranging from 80% to 95% for the five drugs. It indicates that, the anti-HIV activity mainly results from the selective inhibition of HIV replication and less due to toxicity. Their therapeutic index is ranging from 5.9 to 64.8 and far below that of AZT.

**Figure 4.**
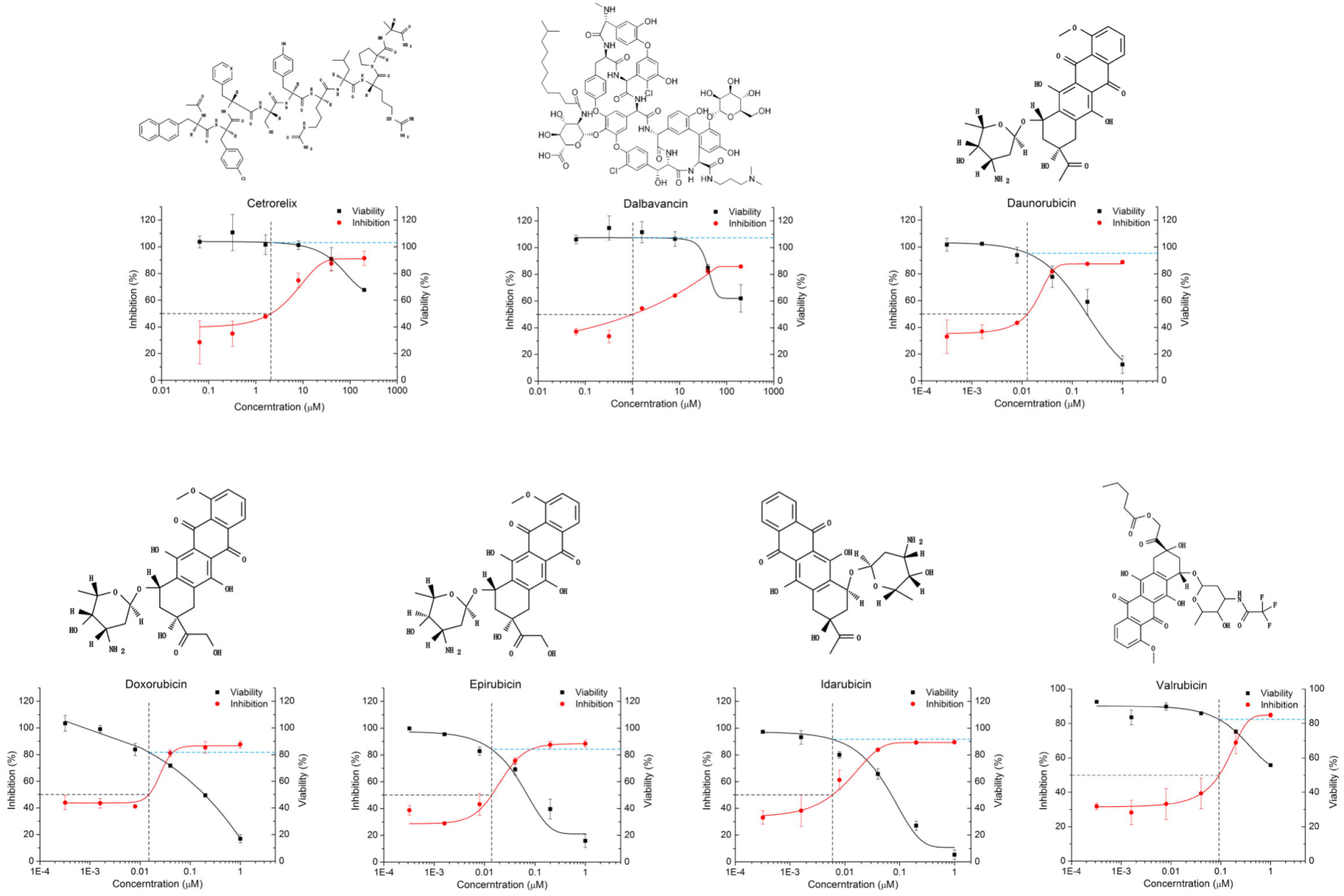
The chemical structures and *in vitro* dose-response curves of the 7 compounds. Each panel contains the structure and curve for one compound. In dose-response curve, the percent inhibition of the compounds on HIV-1 replication in the p24 assay is shown in red circles. And the percent viability in cytotoxicity assays of the compounds on C8166 is shown in filled black squares. With the increase of concentration of the compounds, the percent inhibition is increased but the percent viability of C8166 is decreased. The percent viability at the concentration equal to EC_50_ is indicated as blue dashed line. Data are mean ± s.d. (n=6)

**Table 2.** The anti-HIV-1 activity on HIV-1NL4-3 strain (P24 EC50), and therapeutic index (TI) of the 7 compounds

| **Drug name** | **Original indication** | **EC<sub>50</sub>(μM)**<br>**Mean±SD** | **TI** |
| --- | --- | --- | --- |
| Cetrorelix | For assisted reproduction and the inhibition of premature LH surges | 1.788±0.115 | >105.1 |
| Dalbavancin | For the treatment of acute bacterial infections caused by the Gram-positive pathogens | 1.296±0.186 | >135.1 |
| Daunorubicin | For treatment of leukemia and other neoplasms | 0.016±0.002 | 17.6-36.4 |
| Doxorubicin | For inhibition of disseminated neoplasma like acute leukemia, Hodgkin's disease and so on | 0.012±0.001 | 14.5-18.8 |
| Epirubicin | For adjuvant therapy in patients with breast cancer | 0.011±0.004 | 5.9-23.9 |
| Idarubicin | For treatment of acute myeloid leukemia in adults | 0.003±0.001 | 18.3-40.0 |
| Valrubicin | For treatment of cancer of the bladder | 0.076±0.032 | 18.3-64.8 |
| AZT | For treatment of HIV infections | 0.005±0.004 | >93161 |

## Discussion

The failure of 30 years of HIV vaccine development ^5,35^, as well as the prevalence of drug-resistant HIV^36–38^, emphasizes the need for new, effective and affordable anti-HIV drugs. To decrease the cost and time required for the development of new drugs to treat HIV infection, a world’s first web server Anti-HIV-Predictor was developed for predicting anti-HIV activity of compounds. The accuracy of the web server is more than 93% and AUC is greater than 0.958, which indicates that Anti-HIV-Predictor is powerful enough to discover new anti-HIV agents. Using Anti-HIV-Predictor, 1835 approved drugs were computational screened rapidly. A total of 67 drugs were predicted as anti-HIV compounds. Almost half of the 67 drugs are approved for anti-HIV therapy or with anti-HIV activity. Among the 67 drugs, the drugs with no experimental data for anti-HIV activity were experimentally evaluated in this study. Based on the results of cytotoxicity, syncytia formation and p24 quantification assays, 7 approved drugs (cetrorelix, dalbavancin, daunorubicin, doxorubicin, epirubicin, idarubicin and valrubicin) were identified as new potential anti-HIV agents.

Screening 1835 approved drugs for new anti-HIV drugs through experimental methods is a tedious, expensive and time-consuming process. In this study, the 7 new compounds were rapidly repurposed for anti-HIV therapy from the huge approved drugs library. This process of drug repositioning, which is time and money saved, has benefited from the web server Anti-HIV-Predictor. In silico screen of the approved drugs library using Anti-HIV-Predictor only needs less than one hour. After screening, the predicted anti-HIV compounds can be experimentally evaluated immediately. The rapidity and accuracy of Anti-HIV-Predictor make it powerful for discovery of new anti-HIV agents. In future, we will use Anti-HIV-Predictor to screen other compound database such as TCM Database@Taiwan^39^ and Human Metabolome Database^40,41^ for discovery of new natural product against HIV.

Cetrorelix, a synthetic decapeptide, is used in assisted reproduction to inhibit premature LH surges. The drug works by blocking the action of gonadotropin-releasing hormone (GnRH) upon the pituitary, thus rapidly suppressing the production and action of LH and follicle-stimulating hormone (FSH)^32^. It is administered as 0.25 mg or 3 mg for one subcutaneous injection. The administered dosage is equal to 0.034 - 0.402 μM (0.25 - 3 mg/5L blood) in human blood which is close to the concentration EC_50_ (1.788 μM) for anti-HIV activity. Therefore, administration of cetrorelix as the same for original indication of assisted reproduction may have clinical benefit to HIV-infected patients. Dalbavancin is a novel second-generation lipoglycopeptide antibiotic. It possesses *in vitro* activity against a variety of gram-positive pathogens. Dalbavancin exerts its bactericidal effect by disrupting cell wall biosynthesis^33^. It is administered as 500 mg for one subcutaneous injection. The administered dosage is equal to 55.03 μM (500 mg/5L blood) in human blood which is far higher than the concentration EC_50_ (1.296 μM) for anti-HIV activity. Therefore, administration of dalbavancin as the same for treatment of bacterial infection is very promising to have clinical benefit to HIV-infected patients.

The five anthracycline drugs daunorubicin, doxorubicin, epirubicin, idarubicin and valrubicin identified in this study have potent anti-HIV activity at the nanomolar level. Although they are more toxic than cetrorelix and dalbavancin, their therapeutic index is all more than 10. The therapeutic index of valrubicin is 18.3-64.8, which is the highest among the five drugs. The EC_50_ of idarubicin is 0.003 μM (TI=18.3-40.0), which is best among the five drugs. Idarubicin inhibits HIV-1 replication at the lowest concentration among the five drugs and close to or better than the positive control drug AZT. The five anthracycline drugs are approved for the treatment of lymphomas, leukemias, Hodgkin’s disease, bladder cancer and so on^34^. The HIV-infected patients were more likely to suffer from anal cancer and Hodgkin’s lymphoma^42,43^. HIV-infected patients with cancer are less likely to receive treatment for some cancers than uninfected people, which may affect survival rate ^43,44^. HIV-infected cancer patients are more likely to die from cancer than uninfected cancer patients. Therefore, the five drugs may be applied to treatment of the HIV-infected patients with cancer. These patients may benefit from the five drugs.

Anti-HIV-Predictor predicts anti-HIV activity of compounds based on the benchmark dataset containing active and inactive compounds. The compound with potent anti-HIV activity but less cytotoxicity is expected in the development of anti-HIV drug. Since the cytotoxicity is not taken into account in the current study, some of predicted compounds exhibit high cytotoxicity as shown in Table 1. Therefore, Anti-HIV-Predictor is open to improvement. In future, we will consider the cytotoxicity as important factor in the prediction of anti-HIV activity by integrating the NCI-60 growth inhibition data from NCI Development Therapeutics Program (DTP) (https://dtp.cancer.gov/)^45^. The predicted anti-HIV compounds in the first step will be filtered by cytotoxicity feature. Anti-HIV-Predictor with cytotoxicity filter may results in a compound with high anti-HIV activity but less cytotoxicity.

Treatment of AIDS still faces multiple challenges such as drug resistance and HIV eradication. Development of new, effective and affordable drugs against HIV is urgently needed. Here we firstly developed a world’s first web server Anti-HIV-Predictor for predicting anti-HIV activity of compounds and then applied the server to drug repositioning for anti-HIV therapy. Finally, we repurposed 7 compounds as new anti-HIV agents. The web server and the 7 repurposed anti-HIV agents provided here have an immediate effect on the development of new anti-HIV therapeutics, and should significantly advance current anti-HIV research.

## Materials and methods

### Construction of benchmark dataset

Anti-HIV activity data were downloaded from ChEMBLdb and NCI. In ChEMBLdb, the compound whose target is “human immunodeficiency virus type 1” and with the activity better than 10 μmol/L was considered as active compounds. In NCI, the compound with more than 2 replication experiments and with EC_50_ less than 10 μmol/L was considered as active compounds. And the other compounds with EC_50_ more than 100μmol/L were consider as inactive compounds. Finally, all compounds in the two databases were integrated by removed the conflict and replicated compounds. This procedure yielded 9584 active and 23998 inactive compounds, respectively. The active and inactive datasets were used as benchmark datasets to generate models to predict anti-HIV activity of chemical compounds. The detailed method of constructing benchmark dataset can be found in **Part 1 of Supplemental material.**

### RFW_FP model

Firstly, Relative Frequency-Weighted Fingerprint (RFW_FP) was used to calculate the compound fingerprints. RFW_FP was calculated as follows:

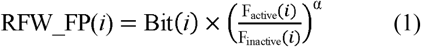

where *i* represents *i*th Daylight fingerprint. In Daylight theory, each compound contains more than one and less than 1024 fingerprints. RFW_FP(*i*) is *i*th relative frequency-weighted fingerprint. Bit(*i*) is calculated by Pybel^46^, a python wrapper of Openbabel^47^. if the compound has *i*th fingerprint, Bit(i) = 1, else Bit(i) = 0. F_active_(*i*) and F_inactive_(*i*) are the frequency of *i*th fingerprint in the active and inactive compounds, respectively. α is the amplifying factor. In this study,α was optimized as 0.5 (**Figure S2**).

Then, the Relative Frequency-Weighted Tanimoto Coefficient (RFW_TC) between two compounds was calculated as follows:

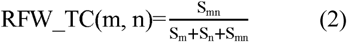

where RFW_TC(m,n) is RFW_TC between two compounds m and n. S_m_ and S_n_ are the sum of RFW_FPs in compound m and n, respectively. S_mn_ is the sum of the common RFW_FPs between two compounds.

Finally, for each query chemical compounds, the maximum RFW_TC between the query and the active dataset (9584 compounds) was calculated. Then the P-value, based on the maximum RFW_TC, was calculated. As the maximum RFW_TC is less than 1.0 and the maximum RFW_TCs of the inactive compounds have a normal distribution (**Figure S3**), we can calculate the P-value as follows:

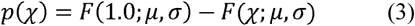

where *p*(χ) is the P-value at the maximum RFW_TC of x; *F*(χ;μ,σ) is the cumulative function of normal distribution. Using the maximum likelihood method (“fitdist” function in R “fitdistrplus” package^48^), we estimated the location parameter μ of 0.461, the scale parameter σ of 0.121.

### SVM model

SVM is a powerful supervised learning algorithm suitable for non-linear classification problems^49^. It is based on the idea of transforming data not linearly separable in feature space to a higher- or infinite-dimensional space where they can be separated linearly by a suitable soft-margin hyperplane^50^. For our binary classification task, we firstly chosen kernel function and then perform a grid search of the penalty parameter C. The Scikit-learn Python wrappers for libsvm27 were used to choose kernel function and explore the hyper-parameter space^51,52^. The best-performing model was selected by plotting receiver operating characteristic (ROC) curve and calculating the area under the curve (AUC). The model with kernel function rbf and the penalty parameter C of 500 performed best (**Figure S4**). The detailed method for the selection of kernel function and the penalty parameter C can be found in **Part 5 of Supplemental material.**

### RF model

The algorithm of random forest is based on the ensemble of a large number of decision trees, where each tree gives a classification and the forest chooses the final classification having the most votes over all the trees in the forest^53^. Random forest, implemented in Scikit-learn^51^, was chosen as classifier with the following settings: (1) Number of trees was set to 900 (n_estimators =900). This parameter was selected by calculating AUC (**Figure S5**). (2) The minimum number of samples to split an internal node was set to 2 (min_samples_split = 2, default setting). (3) The minimum number of samples in newly created leaves was set to 1 (min_samples_leaf = 1, default setting). (4) The number of features to consider when looking for the best split was set to the square root of the number of descriptors(max_features = auto, default setting). (5) The maximum depth of the tree was expanded until all leaves are pure or until leaves contain less than min_samples_split samples (max_depth = none, default setting). (6) Bootstrap samples were used (bootstrap = true, default setting). For further documentation on the random forest implementation in Scikit-learn, the interested reader is referred to the web site (http://scikit-learn.org).

### Performance evaluation

To test performance of Anti-HIV-Predictor, 10 runs of 5-fold cross-validation (CV) method (**Part 2 of Supplemental material**) were used to the three models (RFW_FP model, SVM model and RF model). For each model, the ROC was plotted and the area under the curve (AUC) was calculated. The results of the CV tests were used to calculate the four quality indices: accuracy, precision, recall and F1 score which is defined as the harmonic mean of precision and recall. We used the default statistical definition for these quality indices:

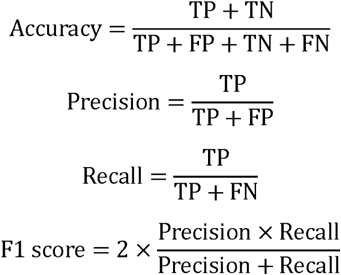

where true positive (TP) and true negative (TN) correspond to correctly predicted anti-HIV compound and non anti-HIV, respectively, false positive (FP) denote non anti-HIV compound predicted as anti-HIV compound, and false negative (FN) denote anti-HIV compound predicted as non anti-HIV compound.

### Compounds, cells and HIV-1 strain

The 28 approved drugs were purchased from CASMART (http://www.casmart.com.cn). C8166 and H9 cell was kindly provided by the AIDS Reagent Project, the UK Medical Research Council (MRC). Cells were maintained in RPMI 1640 medium (Life technology) containing 10% heat-inactivating fetal bovine serum (FBS, Life technology), 100units/mL penicillin (Sigma) and streptomycin (amresco). Laboratory adapted strain HIV-1_NL4-3_ was kindly donated by NIH and propagated in H9 cells. Virus stocks were stored in small aliquots at -70 □.

### Cytotoxicity assays

The cellular toxicity of tested compounds on C8166 was assessed by MTT colorimetric assay^54^. Briefly, 4×10^4^ per well C8166 cells were co-incubated with or without a series diluted test compounds. After 3 days of incubation at 37 □, 5% CO_2_, the cell viability was determined by using MTT. Afterward, the 50% cytotoxicity concentration (CC_50_) was calculated. AZT was used as a positive control.

### Inhibition of syncytia formation

The inhibitory effect of samples on acuteHIV-1_NL4-3_ infection was measured by the syncytia formation assay as described previously^55^. In the presence or absence of various concentrations of compounds, 4×10^5^/ml C8166 cells were infected with HIV-1_NL4-3_ at a multiplicity of infection (MOI) of 0.03, and cultured in 96-well plates at 37 □ in 5% CO_2_ for 3 days. AZT was used as a positive control. After post-infection for 3 days, cytopathic effect (CPE) was measured by counting the number of syncytia in each well of 96-well plates under an inverted microscope (10×) (Nicon ECLIPSE TS100). The inhibitory percentage of syncytia formation was calculated by the percentage of syncytia number in treated sample compared to that in infected control. 50% effective concentration (EC_50_) was calculated. Therapeutic index (TI) was calculated by the ratio of CC_50_/EC_50_.

### Inhibition of HIV-1 p24 antigen level in acute infection

For the compounds with TI value greater than 10, the in vitro inhibitory effect of these compounds on HIV-1 replication was further evaluated by quantification of p24 expression. Briefly, 4×10^5^/ml C8166 cells were infected with HIV-1_NL4-3_ for 2 hours to allow for viral absorption. It was then washed three times with PBS to remove unadsorbed virus. The cells were plated at 4×10^4^ cells/well with or without various concentrations of compounds and incubated in a CO_2_ incubator at 37 °C with for 72 hours. Supernatants were collected and virus was lysed with 0.5% triton X100. HIV-1 p24 was determined with an in-house ELISA assay described previously^31^. The inhibitory percentage of p24 antigen production was calculated by the OD_490/630_ value of compound-treated culture compared to that in infected control culture and EC_50_ were calculated.

## Acknowledgements

This work was supported by the National Basic Research Program of China (Grant No. 2013CB835100), the Key Scientific and Technological Program of China (2014ZX10005-002-006), the National Natural Science Foundation of China (Grant Nos. 31401142, 31401137, 81471620, 81660612), the Instruments Function Deployment Foundation of CAS (2014gk01) and the Applied Basic Research Foundation of Yunnan Province (2014FB181; 2015FB182; P0120150150).

## Author contribution

S-X D, Y-T Z and J-F H conceived and designed the research. S-X D, H C,W-X L, Y-C G, J-Q L, J-J Z, Q W, H-J L, B-W C, Y-D G performed data analysis. H C performed the in vitro experiments of anti-HIV activity and cytotoxicity assays. S-X D, H C, W-X L, G-H L, Y-T Z and J-F H wrote or contributed to the writing of the manuscript. All authors reviewed the manuscript.

